# Collateral sensitivity networks reveal evolutionary instability and novel treatment strategies in ALK mutated non-small cell lung cancer

**DOI:** 10.1101/075846

**Authors:** Andrew Dhawan, Daniel Nichol, Fumi Kinose, Mohamed E. Abazeed, Andriy Marusyk, Eric B. Haura, Jacob G. Scott

## Abstract

Drug resistance remains an elusive problem in cancer therapy, particularly with novel targeted therapy approaches. Much work is currently focused upon the development of an increasing arsenal of targeted therapies, towards oncogenic driver genes such as ALK-EML4, to overcome the inevitable resistance that develops as therapies are continued over time. The current clinical paradigm after failure of first line ALK TKI is to administer another drug in the same class. As to which drug however, the answer is uncertain, as clinical evidence is lacking. To address this shortcoming, we evolved resistance in an ALK rearranged non-small cell lung cancer line (H3122) to a panel of 4 ALK tyrosine kinase inhibitors used in clinic, and performed a collateral sensitivity analysis to each of the other drugs. We found that all of the ALK inhibitor resistant cell lines displayed a significant cross-resistance to all other ALK inhibitors. To test for the stability of the resistance phenotypes, we evaluated the ALK-inhibitor sensitivities after drug holidays of varying length (1, 3, 7, 14, and 21 days). We found the resistance patterns to be stochastic and dynamic, with few conserved patterns. This unpredictability led us to an expanded search for treatment options for resistant cells. In this expansion, we tested a panel of 6 more anti-cancer agents for collateral sensitivity among the resistant cells, uncovering a multitude of possibilities for further treatment, including cross-sensitivity to several standard cytotoxic therapies as well as the HSP-90 inhibitors. Taken together, these results imply that resistance to targeted therapy in non-small cell lung cancer is truly a moving target; but also one where there are many opportunities to re-establish sensitivities where there was once resistance.

## 1 Introduction

Drug resistance in cancer is fundamentally an evolutionary problem [1]. Tumour cells, within the varying microenvironment of a patient, are subjected to the selective pressures of the drugs to which they are exposed, and respond in a manner governed by Darwinian dynamics [2]. As a consequence of these stochastic evolutionary dynamics, the resultant population of drug-resistant cancer cells may display features of cross-resistance, or conversely, collateral sensitivity, to other chemotherapeutic agents; a knowledge of which, may be used to guide further therapy. Collateral sensitivity is sensitivity towards a second drug which occurs after the evolution of resistance to a first drug, when the resistant state causes a vulnerability to another drug that was not previously present [3]. Clinically, a case of collateral sensitivity by resistance mutations, or sensitization to a second drug in a state of resistance to a first-line therapy, has been shown in a patient with ALK-positive NSCLC [4]. Further, the utility of a broader knowledge of collateral sensitivity to panels of drugs has been shown in E. coli and lymphoma alike [3, 5].

In addition to the concept of collateral sensitivity, drug holidays, or metronomic therapy, have also been proposed to limit the development of resistance (or analagously, extend efficacy) in cancer treatment [6, 7, 8]. Upon removal of the selection pressure (therapy), it is no longer necessarily advantageous to possess the changes associated with resistance. If the changes associate with resistance come at a significant cost then, these traits may be selected out of the population in the absence of drug. The molecular basis of the efficacy of drug holidays, in a general sense, is thought to be due to reversible non-mutational mechanisms [6]. Little is known, however, about the length of drug holiday necessary for the outgrowth of the original, sensitive populations, and this is likely highly variable from cancer to cancer, and from patient to patient, though indeed, small-scale clinical studies have shown benefit with this technique [6, 9]. Recent work in this area has also shown that adaptive therapy, or the careful titration of therapy to maintain a stable tumour burden, but not the eradication, to prevent resistance and ultimate therapeutic failure, has significant clinical promise [10, 2].

In this work, we combine the ideas of drug holidays and collateral sensitivity to develop strategies of overcoming TKI resistance in ALK-positive NSCLC. First described in 2007, rearrangements in the anaplastic lymphoma kinase (ALK) and the echinoderm microtubule-associated protein-like 4 (EML4) genes have been found to drive approximately 5-10% of all non-small cell lung cancer (NSCLC) cases, disproportionately affecting younger, generally non- or light smoking, patients [11, 12, 13]. Clinically, targeted therapies inhibiting the kinase activity of ALK have proven to be efficacious, significantly extending progression-free survival compared to standard therapies. These trials have led to the ALK inhibitor Crizotinib to be the first-line standard of care for metastatic tumours driven by this oncogene [14, 15]. After widespread use began however, reports of resistance to ALK inhibition quickly emerged, and it has since become apparent that within one year of starting such therapy, resistance almost inevitably emerges [16, 17, 18]. The clinical question that arises thereafter is how to proceed with therapy, and current National Comprehensive Cancer Network (NCCN) guidelines suggest that for a symptomatic patient, a second-line agent of the ALK-TKI class should be used, such as Ceritinib or Alectinib [14]. Reportedly, these next generation ALK TKIs can overcome resistance to Crizotinib [19], but there is little guidance as to how to choose the most efficacious second line therapy. There are, however, alternative strategies not represented in these guidelines that may provide avenues for further treatment. One such strategy to combat therapeutic resistance, with reports of possible efficacy in lymphoma [5], involves exploiting collateral sensitivity.

In this work we derive what is, to our knowledge, the first collateral sensitivity network using targeted therapies in any solid cancer, and extend the analysis to the period after the drug is removed. We posit that drug holidays are an important, yet overlooked aspect of clinical treatment. Typically, when anti-cancer therapies are switched, after progression say, there is a latency period in which the patient is not receiving any form of anti-cancer therapy, on the order of weeks to months. This latency period, or holiday, has not been considered in previous experimental or theoretical models. This led us to investigate the evolutionary dynamics of collateral sensitivities in drug resistant cell lines after having undergone various lengths of drug holidays. In particular, we examine the structure of the collateral sensitivity network for the available ALK TKI in an ALKmut NSCLC line. We further investigate the effect of drug holidays on this structure, and the potential for novel cycling regimens using a panel of clinically relevant therapeutic agents. We show that evolution of resistance to a particular ALK-TKI leads to a substantial cross-resistance to other ALK-TKIs, but this varies significantly over time with few conserved motifs. Further, we show that an expanded panel of drugs (full panel described in Table 1) provides a significant range of novel possible treatment regimens, including multi-drug cycles, that may be tested clinically [20, 21].

**Table 1:** Panel of drugs used in study, their abbreviations, and classification.

| Drug name | Abbreviation | Class |
| --- | --- | --- |
| Ceritinib | Cerit | ALK-TKI |
| Alectinib | Alec | ALK-TKI |
| Lorlatinib | Lorl | ALK-TKI |
| Crizotinib | Criz | ALK-TKI |
| Paclitaxel | Pacl | Taxane |
| Ganetespib | Gane | Hsp90 Inhibitor |
| IPI504 | IPI504 | Hsp90 Inhibitor |
| AUY022 | AUY922 | Hsp90 Inhibitor |
| Pemetrexed | Pem | Folate Antimetabolite |
| Etoposide | Etop | Topoisomerase Inhibitor |

## 2 Results

### 2.1 Significant cross-resistance among ALK-TKI resistant cells

Evolution of resistance to ALK inhibitors (Crizotinib, Alectinib, Lorlatinib, and Ceritinib), was induced over a 16 week period beginning with wild type H3122 ALK-positive NSCLC cell line, and cell lines were subjected to a collateral sensitivity analysis, as described in Methods. From the obtained EC50 values in the dose-response experiments for each of the resistant cell lines, we compare the collateral resistances and sensitivities between the cell lines. The data is presented as a heat map of fold changes of the EC50 values compared to the values for the treatment naive parental cells (Figure 1). Contrary to the accepted clinical guidelines, none of the 4 resistant cell lines obtained displayed collateral sensitivity to any other ALK TKI at the time of maximum drug exposure. Further, with the exception of Lorlatinib treatment following Alectinib resistance, which was neutral, cross-resistance was observed among all four resistant cell lines.

**Figure 1:**
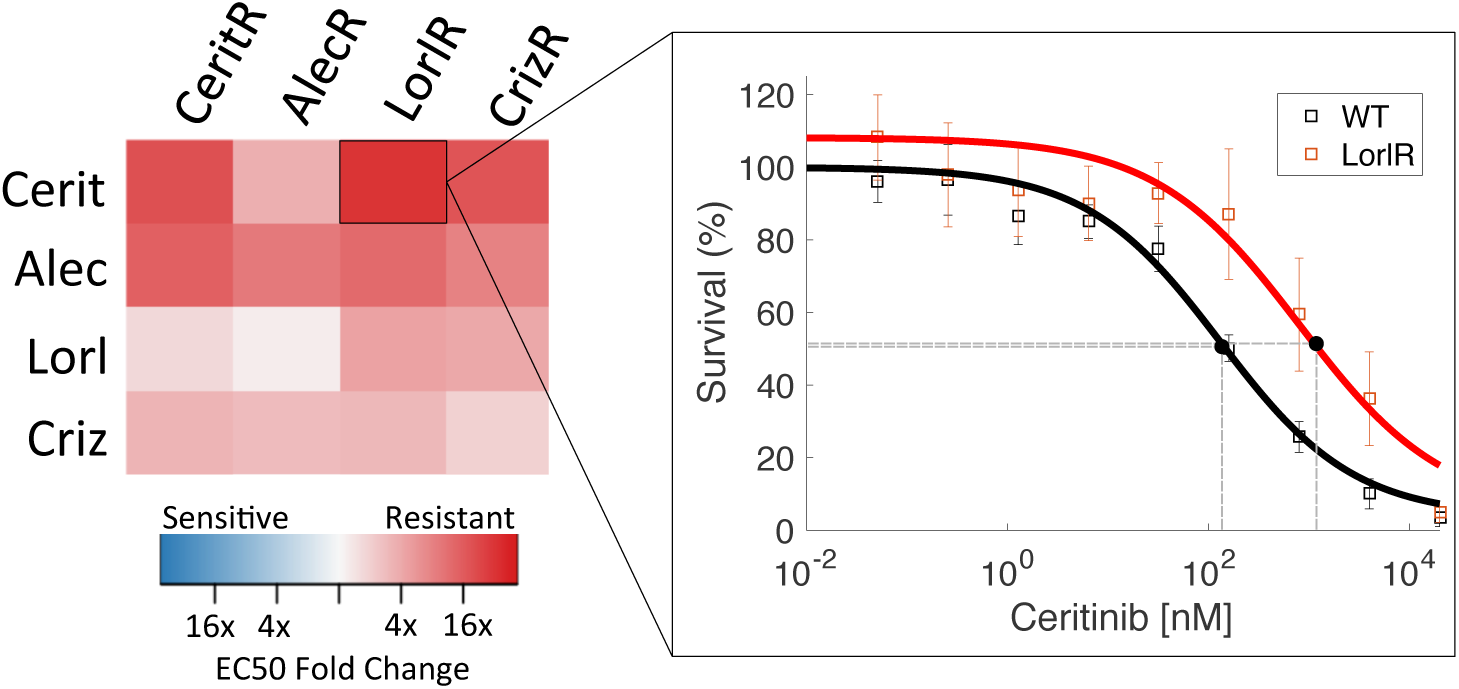
Left: Collateral sensitivity matrix of fold change of EC50 for resistant cell lines (columns) as treated by the panel of ALK-TKIs (rows). All sequences of therapy resulted in cross-resistance except Alectinib followed by Lorlatinib, which was neutral. Right: Pop-out figure shows example of EC50 comparison in case of collateral resistance of Lorlatinib resistant cells treated with Ceritinib, as compared to wild type (WT). Experimental data (markers) and model fit (solid lines) are shown.

### 2.2 Drug holidays stochastically induce collateral sensitivity between ALK TKIs with few conserved motifs

In the clinical setting, drug holidays have been suggested as a strategy to overcome therapeutic resistance, as resistance may not be preserved throughout time. Furthermore, there is often a substantial time period in which no drug is given, after the administration of the first drug, and prior to the administration of the next. This drug holiday may affect the efficacy of the drug sequencing protocol, but is often neglected in experimental and theoretical studies of resistance alike. It is therefore of critical importance to assay the stability of possible sequencing regimens not only among cell lines in which resistance has been derived, but also in which drugs have been stopped for a period of time, to simulate clinically-relevant situations. To address this, we assayed the four resistant cell lines for five drug holiday periods: 1 day, 3 days, 7 days, 14 days, and 21 days.

After assaying each of the cell lines for drug response, we construct the temporal collateral sensitivity matrices (Figures 2A - E, left) and derive the resultant sensitivity networks in Figures 2A - E, right. For details on graph construction and associated code, see Methods. We find that there are patterns that change particularly quickly, such as the collateral sensitivity to Lorlatinib in Ceritinib resitant cells, appearing on the first day of holiday, then disappearing on day 3, re-appearing on day 7, disappearing on day 14, and again re-appearing on day 21. Likewise, there are also stable patterns that are highly conserved, though few in number, such as the cross-resistance of Lorlatinib resistant cells towards Ceritinib. Further implication of these experimental results indicates that Lorlatinib might be a poor choice for first-line therapy, as the cross-resistance it generates is significant and stable through drug holidays. Lorlatinib, as indicated by these results, for this cell line, may be a better choice in the second-line, as many resistant lines are sensitized towards it after a drug holiday.

**Figure 2:**
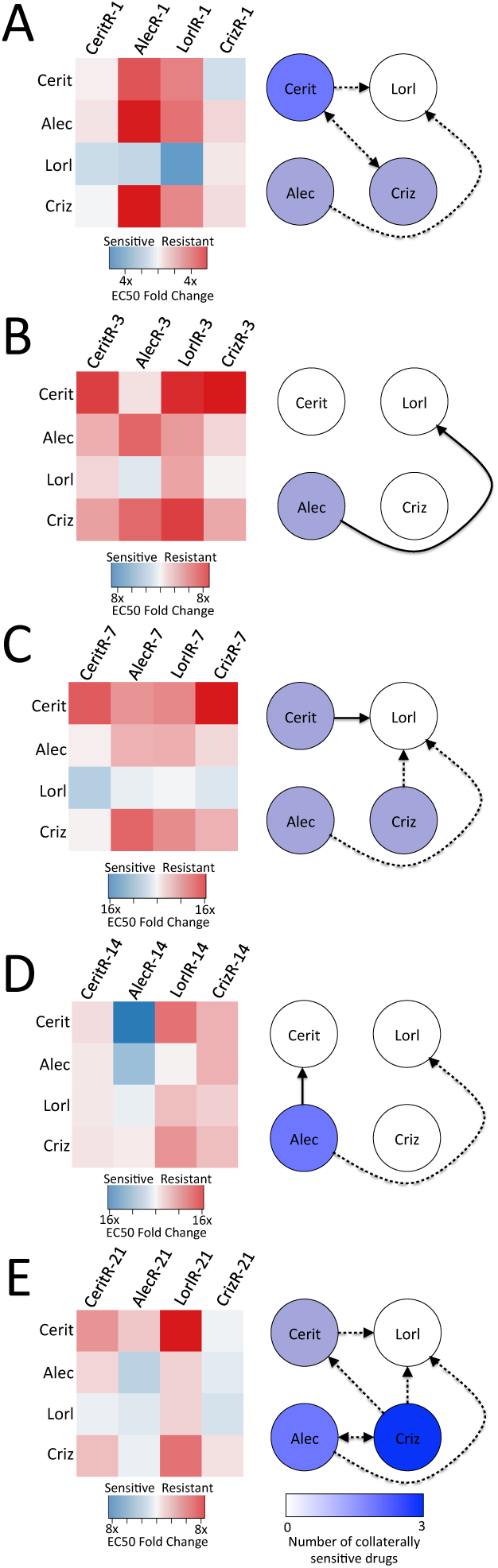
Left: Collateral sensitivity matrices depicting fold change of EC50 for resistant cell lines, during the therapy holiday lasting for 1 day (A), 3 days (B), 7 days (C), 14 days (D), and 21 days (E), as treated with the panel of ALK-TKIs (rows). Right: Collateral sensitivity networks depicted as directed graphs for cells during therapy holiday for 1 day (A), 3 days (B), 7 days (C), 14 days (D), and 21 days (E). Each named node represents the resistant population to that drug; an edge exists from node *i* to node *j* if cells resistant to drug *i* are sensitive to drug *j*. The colour of each node represents the number of drugs to which the cell line represented by that node is sensitive (i.e. the number of out edges). Dotted arrows indicate those in which 95% confidence intervals for wild-type EC50 overlapped with that of cell line tested.

Another implication of these experimental results lies in the design of metronomic chemotherapy schedules. That is, there appears to be a dynamic sensitization of Alectinib resistant cells to Alectinib, appearing only after 14 days of drug holiday, and maintained after 21 days of holiday as well. This suggests the design of a schedule in which one treats in regular intervals, but with a 14 or 21 day holiday in between cycles, as in metronomic chemotherapy.

Finally, we note that variation in these graphs highlights significant temporal instability in collateral sensitivities over time. These dynamics have largely not been taken into account in previous studies of collateral sensitivity and resistance, in both the fields of microbiology and oncology [3, 22, 23, 24, 25]. Based on these experiments, we pose that this is a critical phenomenon that must be accounted for when designing treatment schedules, and may have significant implications for previously obtained results of collateral sensitivity.

### 2.3 Expanded drug panel provides opportunities for significant collateral sensitivity to ALK TKIs

After finding significant cross resistance between all ALK TKIs, we expanded our search by testing the sensitivities to clinically relevant chemotherapeutic agents and heat shock protein inhibitors. While clinical trials have shown the superiority of targeted therapy in the first-line setting, there is currently little to no data about the relative efficacy of non-targeted chemotherapy in the setting of therapeutic resistance. This avenue presents significant opportunity for intervention in the case of resistance to the ALK TKIs, as we have found that there are a number of drugs, not in the class of TKIs, to which a number of the resistant cell lines are collaterally sensitive, as depicted in Figures 3A and B. We find that the cell lines resistant to the first-line therapy of TKIs can often by sensitized towards non-TKIs in the resistant state. In order to evaluate the various drugs in a general sense, we may consider the number of resistant cell lines that display collateral sensitivity or cross-resistance towards a given drug, which we plot in Figure 3B. While not an exhaustive, or mechanistic explanation, this may be used to consider the potential for the use of a given therapy as a second-line agent in a probabilistic sense, though we recognize that in a clinical setting this probability is adjusted (conditioned) by knowledge of the first-line agent. Ranking drugs by the probability that a given resistant cell line is collaterally sensitized towards them, as is done in Figures 3C and D, we observe that Etoposide and Pemetrexed are the optimal choices for second line therapy, as the greatest number of resistant cell lines are sensitized towards them. Further, ranking by number of drugs to which cross-resistance exists in Figure 3D, we observe that the HSP-90 inhibitors, Ganetespib and IPI504, are the poorest choices for second-line therapy after the emergence of resistance, as they carry the greatest burden of cross-resistance.

**Figure 3:**
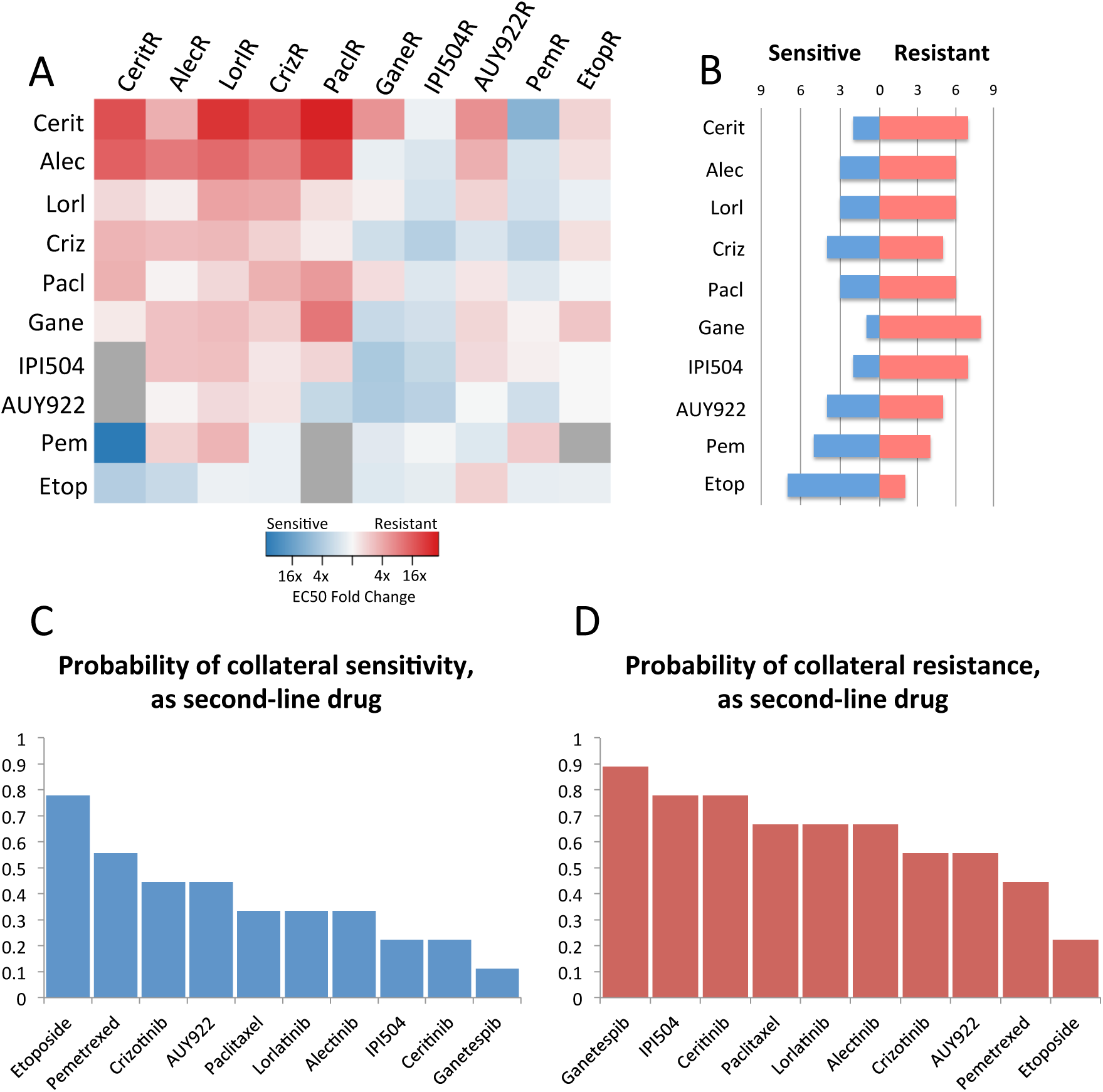
(A): Collateral sensitivity matrix of fold change of EC50 for resistant cell lines (columns) as treated by the panel of anti-cancer therapies (rows). Grey boxes are indeterminate due to significant resistance to drug. (B): Bar graphs depicting the number of collaterally sensitive or resistant cell lines to each drug. (C): Ranked probabilities of collateral sensitivity of each drug to the resistant cell lines, when the drug is used as second-line therapy. (D): Ranked probabilities of cross-resistance of each drug to resistant cell lines, when the drug is used as second-line therapy.

Beyond assaying efficacy of these agents in the second line setting, or as drugs to prime a cancer to be more sensitive to a targeted agent, this expanded panel of collateral sensitivities and resistances can be taken advantage of to design drug cycling protocols. Because of the patterns of collateral sensitivity observed in Figure 3A, a connected graph of 10 nodes, each node representing a resistant cell line, can be generated (Figure 4A). From this, determining drug cycling protocols consists of identifying graph cycles, which represent sequences of drugs wherein sensitization towards the drugs repeats, and resistance to the final drug results in sensitivity to the first. Counting the number of cycles, and organizing this quantity by length of cycle, we find there are a combined total of 84 unique cycles that can be chosen of length 2-7 (see Figure 4B). We highlight an example of a length 4 cycle by the red arrows of Figure 4A (Crizotinib, Etoposide, Paclitaxel, AUY922). These cycles represent theoretical protocols to test further, though whether they act stably through time, or indeed can be repeated, we leave for future work.

**Figure 4:**
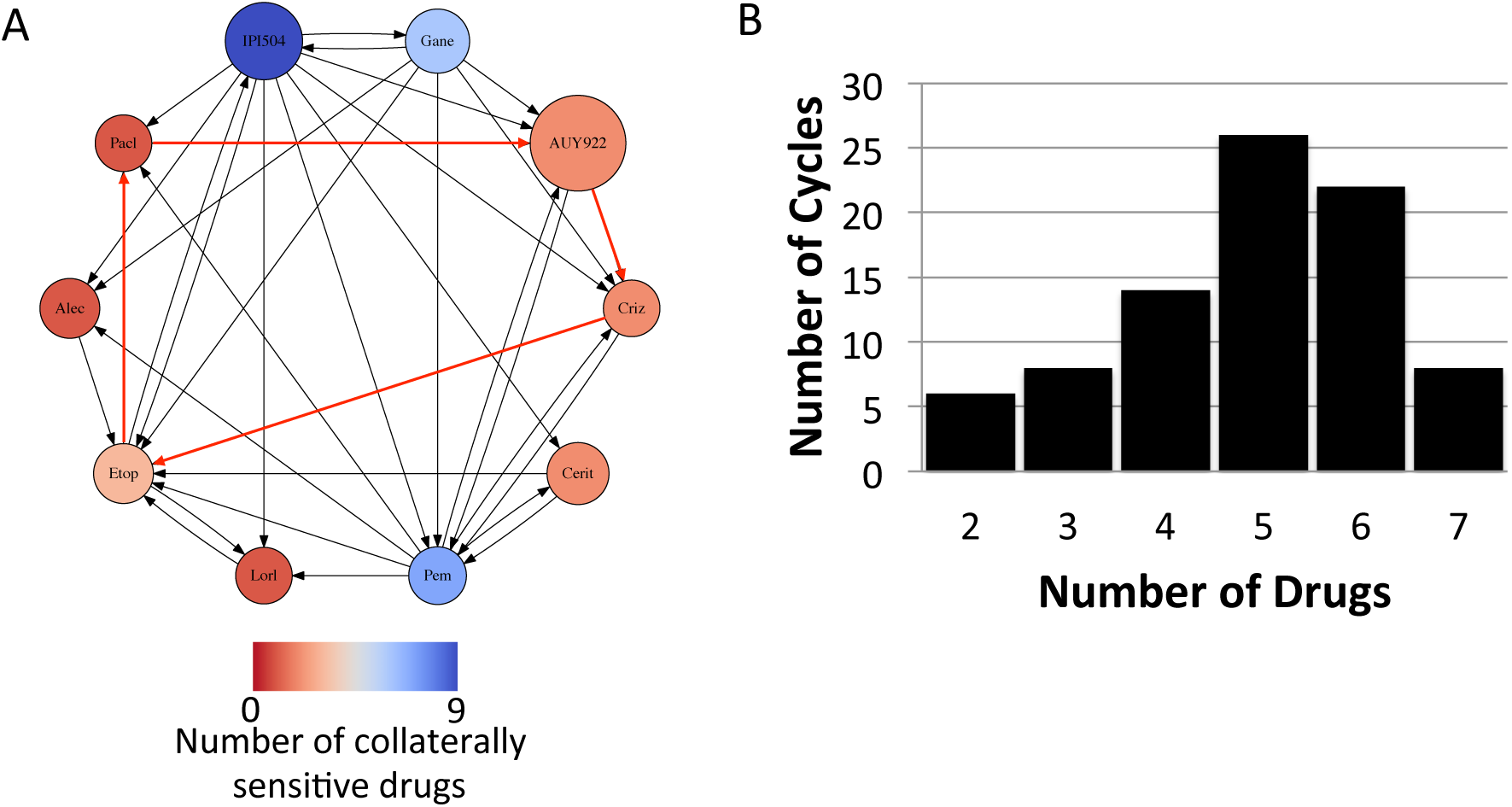
(A): Collateral sensitivity network depicted as a directed graph, with edges pointing from nodes at which resistance to a particular drug has developed to a node for which sensitivity of the drug has increased. Arrows in red indicate an example of a cycling regimen of 4 drugs. (B): Graph of number of drug cycles of given length in the graph presented in (A).

## 3 Discussion

In this work, four cell lines individually resistant to a panel of first-line therapies for ALK-positive NSCLC were studied to determine patterns of resistance and sensitivity, from which to infer second-line treatments. From the data collected, we have constructed a collateral sensitivity ‘map’ for cell lines resistant to the ALK-TKIs. From this we immediately observe that there is significant cross-resistance that evolves under treatment from the first ALK-TKI to the others, suggesting that using any of these drugs as a second-line agent would not provide optimal therapeutic benefit. This differs from previous results, which offer observations of collateral sensitivity among drugs of the same class, as in the case of BCR-ABL driven leukaemia treated with tyrosine kinase inhibitors [5]. Following this, we assay the stability of this cross-resistance after a drug holiday of 1, 3, 7, 14, or 21 days and find significant variation in collateral sensitivity network structure as time progresses, with collateral sensitivities arising transiently and unpredictably with few conserved motifs. In contrast, related work had not considered this possibility of temporal variation after removal of therapy, and instead found particular genetic changes conserved among collaterally sensitive clones, which are assumed to be temporally stable [5]. A critical implication of these results is that even after relatively short drug holidays, conserved motifs are rare among the collateral sensitivity networks constructed. This suggests that strategies involving drug holidays and collateral sensitivities offer promise as collateral sensitivity does arise, but we must carefully consider the dynamics if we are to use these strategies in the clinical setting.

Given the significant cross-resistance which arose in our evolved cell lines, a larger panel of drugs was tested for collateral sensitivity, with encouraging results. Though it has been shown that clinically these drugs are inferior to ALK-TKIs as first-line therapy in patients with NSCLC harboring the ALK rearrangement, we find that there may be utility in considering these drugs after initial therapeutic resistance to ALK-TKIs occurs, with many opportunities for collateral sensitivity (Figure 4). Further, in cases with greater degrees of collateral sensitivity, such as with this expanded panel, as has been reported in this work and many others, the potential for steering or drug cycling regimens exists [3, 5, 26, 27, 28], which would allow us to sequence targeted and non-targeted agents together to regain sensitivity where there was once resistance. In this manner, resistance towards a drug sensitizes for the next drug, and so on, until the drugs of the cycle repeat, and we have shown in Figure 4D that there are a large number of possibilities for these cycling regimens, even for the relatively small panel of drugs tested.

Overall, the number of possible drug cycling regimens in Figure 4D provides hope that expanding the arsenal of drugs will give novel avenues for therapy, and opportunities to reinstate sensitivity, not accounted for in current clinical practices. However, we note that as we have shown in Figure 2, collateral sensitivity is highly dynamic, and truly represents a ‘moving target.’ In this sense, we note that any chosen drug cycling regimen should show stability over time, through drug holidays of varying lengths, as may occur when the drugs of the cycles are being switched. In recent work with *E. coli*, drug cycling protocols based on collateral sensitivity are highlighted, but this distinction based on temporal instability was not raised [3] in regards to timing of the second (and subsequent) therapies. Further, the observation that the collateral sensitivity networks change over time in the absence of drug raises the question as to the stability and repeatability of the generation of the collateral sensitivity matrix itself.

Answering this question will require many biological replicates of each evolution experiment to determine how robust the development of these networks truly is, in solid tumors.

Regardless of the caveats, the observations and method of understanding drug sequencing presented here represent a novel way to utilize existing drugs to regain the upper hand in the clinics against drug resistance, without the need for costly new drugs.

## 4 Methods

### 4.1 Experimental methods

#### 4.1.1 Reagents

Crizotinib, Alectinib, Ceritinib, Paclitaxel, Pemetrexed, Etoposide, Luminespib (AUY-922) and Lorlatinib (PF-06463922) were purchased from ChemieTek (Indianapolis, IN). Ganetispib was purchased from Selleckchem (Houston, TX). Retaspimycin hydrochloride (IPI-504) was purchased from Apexbio (Houston, TX).

#### 4.1.2 Cell culture

H3122 cells were cultured in RPMI 1640 supplemented with 10% FBS (both from Invitrogen, Carlsbad, CA). The cell line was authenticated by STR analysis and tested negative for mycoplasma (PlasmoTest, InvivoGen, San Diego, CA). Cell viability was determined by CellTiter-Glo® Luminescent Cell Viability Assay (Promega, Madison, WI) from cells seeded at 1000 cell/well in 384-well assay plates and drug dosed for 72 hours.

#### 4.1.3 Generation of drug resistance

A clonal line derived from H3122 was used to generate drug resistant cell lines to the 10 drugs listed above, as depicted in Figure 5A, and following the method outlined by Katayama et al. in [29]. Briefly, CellTiter-Glo® was used to determine a baseline IC50 value in the clonal H3122 cell line for each of the 10 drugs. H3122 cells were then seeded at 70% confluence and drugs were added at a starting concentration of 20% of the determined IC50 value. Media/drug was changed every 3 days and cells were passaged once they reached 90% confluency. After two passages, the drug concentration was increased by 2.5-5x fold. This sequential, increased drug exposure was continued for 4-6 months.

**Figure 5:**
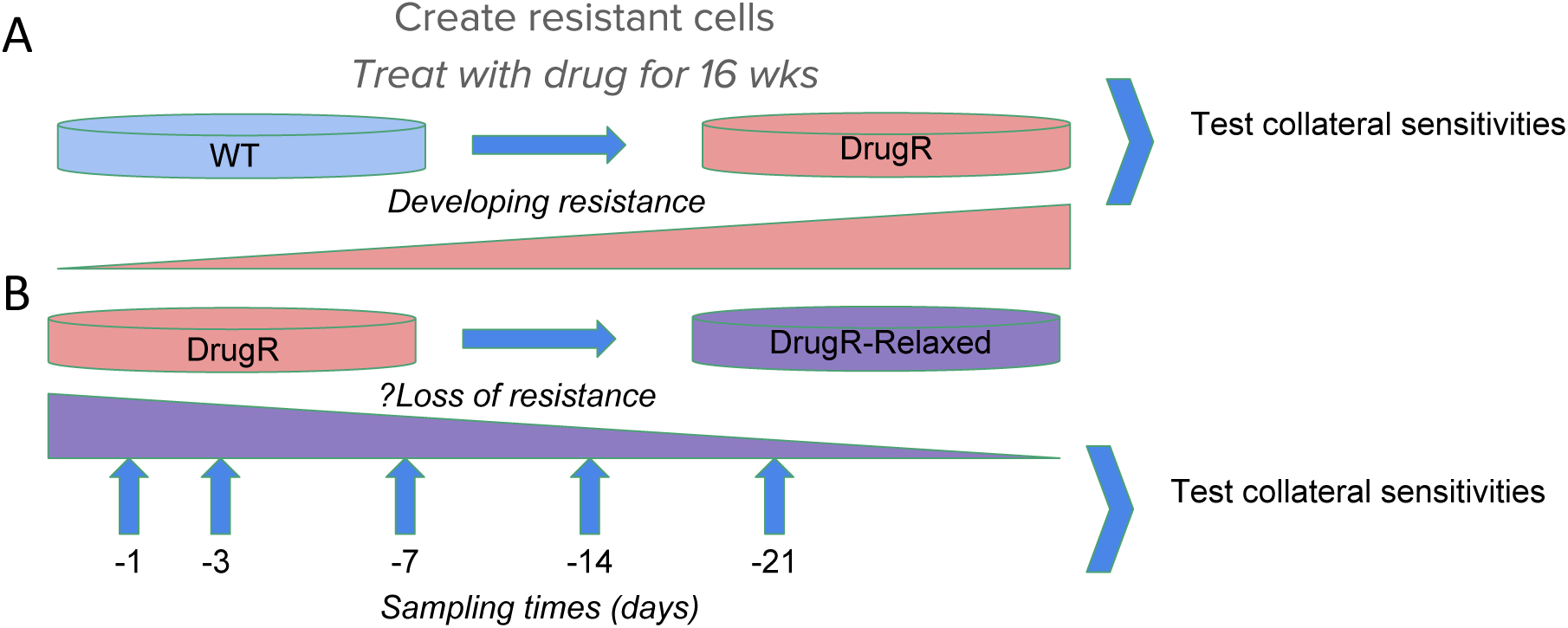
Experimental design diagram. Panel (A) depicts the evolution of resistance by exposure for 16 weeks to drug to create a drug-resistant population (DrugR) and in (B) the population is removed from drug (‘relaxed’).

#### 4.1.4 Drug relaxation and collateral sensitivity assays

H3122 cell lines resistant to tyrosine kinase inhibitors (TKI, Crizotinib, Alectinib, Ceritinib and Lorlatinib) were used for drug relaxation assays. The four TKI drug resistant cell lines were cultured in the presence or absence of the specific drug used in generating the cell line for 1, 3, 7, 14 and 21 days (Figure 5B). To assay cell viability at each of these time points, survival was determined against the 10 drug library using CellTiter-Glo® as described above.

### 4.2 Mathematical analysis

Experimentally, dose-response curves for each cell line to each drug were obtained at varying concentration levels. To extract meaningful comparison from these sets of curves, these were fit using non-linear optimization in R to the following mathematical model. Key model parameters are EC50, which for the purposes of this experiment is defined as the concentration of the drug at which the half-maximal effect of cell kill was observed, and the hill coefficient, *n*. We then model the survival *S*(*d*) after the administration of a particular drug dose *d* as:

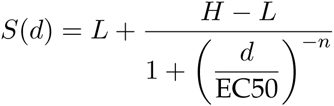

where *n* is the Hill coefficient, *H* is the maximum survival proportion observed, and *L* is the minimal survival proportion observed. In certain cases of data fitting using this procedure, results obtained for the EC50 are reported as indeterminate. These curves represented cases in which cellular resistance was so significant that growth continued in the presence of drug, resulting in a survival function that was not of Hill-type, and could therefore not be meaningfully fit to obtain an EC50.

We note that while there is no perfect measure of drug resistance or sensitivity, the EC50 was chosen in this experiment for the primary purpose of comparability with existing literature, and because of the strong degree of agreement of this model to experimental data, underscoring its utility in comparing dose-response curves. Importantly, we note that there are cases in which the EC50 may mislead as to whether a cell line is resistant or sensitive. For instance, there is the possibility that the reported EC50 for a cell line is lower than the wild type case, but the value of *L* is much greater, meaning that a much smaller proportion of cells were actually eliminated, even at maximal drug concentrations.

### 4.3 Detection of cycles

Once the graph defined by the collateral sensitivity network was reconstructed, cycles were counted from this graph. This was done computationally in Python, with code as provided in the supplementary files. The algorithm relies upon a depth-first search to traverse the graph, iterating through the graph, not revisiting any nodes it has already visited in that iteration of the search. The ‘depth’ of the search is restricted to be the cycle length. The search is then iterated over the graph, and if the starting node is reached at exactly the depth of the desired cycle length, that path is saved as a cycle. We note that this method, by its definition, counts every cycle of length *n*, *n* times, so we divide by this when recording the number of unique cycles.

## 5 Acknowledgments

The authors would like to thank Dr. Nima Sharifi and Mr. Artem Kaznatcheev for helpful discussions and commentary on the manuscript during preparation.

